# HD-Zip II transcription factors control distal stem cell fate in Arabidopsis roots by linking auxin signaling to the FEZ/SOMBRERO pathway

**DOI:** 10.1101/2023.12.06.570393

**Authors:** Marco Possenti, Giovanna Sessa, Altea Alfè, Luana Turchi, Valentino Ruzza, Massimiliano Sassi, Giorgio Morelli, Ida Ruberti

## Abstract

In multicellular organisms, specialized tissues are generated by specific populations of stem cells through cycles of asymmetric cell divisions, where one daughter undergoes differentiation and the other maintains proliferative properties. In *Arabidopsis thaliana* roots, the columella - a gravity-sensing tissue which protects and defines the position of the stem cell niche - represents a typical example of a tissue whose organization is exclusively determined by the balance between proliferation and differentiation. The columella derives from a single layer of stem cells through a binary cell fate switch that is precisely controlled by multiple, independent regulatory inputs. Here, we show that the HD-Zip II transcription factors HAT3, ATHB4, and AHTB2, redundantly regulate columella stem cell fate and patterning in the Arabidopsis root. The HD-Zip II transcription factors promote columella stem cell proliferation by acting as effectors of the FEZ/SMB circuit and, at the same time, by interfering with auxin signaling to counteract hormone-induced differentiation. Overall, our work shows that HD-Zip II connect two opposing parallel inputs to fine tune the balance between proliferation and differentiation in columella stem cells.

**Summary statement:** HD-Zip II transcription factors redundantly regulate columella stem cells activity by counteracting auxin-mediated differentiation downstream of the FEZ/SMB proliferative input

## Introduction

The balance between cell proliferation and differentiation, which is key for the generation of specialized tissues in multicellular organisms, is governed by asymmetric cell divisions (ACD). When a cell divides asymmetrically, it generates two daughter cells with different fates: one retains the self-renewal properties of the mother, whereas the other acquires a specific cell fate and is committed to differentiate (Pierre-Jerome et al., 2018; Pillitteri et al., 2016). In plants, where the presence of a cell wall limits the movement of cells within the tissues, the coordination between cell proliferation and differentiation in the shoot and root apical meristem (SAM and RAM, respectively) is essential to ensure proper developmental processes throughout the plant’s life span (De Smet and Beeckman, 2011).

Because of its stereotypical organization, the RAM of *Arabidopsis thaliana* represents an ideal system to study the coordination between proliferation and differentiation in plant development. The stem cell niche (SCN) in the *Arabidopsis* RAM is centered around a group of slowly replicating cells, the quiescent center (QC) which functions as stem cell organizer (Scheres, 2007). The QC is surrounded by two different pools of stem cells (also known as initials) that produce daughters that give rise to all the tissues of the root (Dolan et al., 1993; Scheres, 2007). The daughters located at the proximal and lateral sides of the QC form a transit-amplifying population (the proximal RAM) that undergoes several rounds of divisions before differentiating into specialized cells. By contrast, the daughters of columella stem cells (CSC), which are located distally to the QC, directly differentiate into columella cells (CC) by accumulating starch granules in amyloplasts for gravity sensing (Scheres, 2007). Because CC ultimately detach from the root cap, the coordination of CSC proliferation and daughter cell differentiation must be tightly controlled to maintain a constant number of CC tiers, thus a correct positioning of the SCN within the RAM (Scheres, 2007).

A number of independent yet intertwined pathways regulate the maintenance of distal stem cells by controlling the activity of the QC, the proliferation of CSC, and the differentiation of daughter cells (Pardal and Heidstra, 2021; Shimotohno and Scheres, 2019). The homeobox gene *WUSCHEL-RELATED HOMEOBOX 5* (*WOX5*), has a central role in regulation of distal stem cells. WOX5 is expressed in the QC where it acts cell-autonomously to suppress cell divisions by excluding CYCLIN D3;3 activity from the QC (Forzani et al., 2014). On the other hand, WOX5 suppresses the differentiation of CSC non-cell-autonomously by repressing the expression of *CYCLING DOF FACTOR 4* (*CDF4*) via chromatin modifications upon interaction with TOPLESS (TPL) corepressor (Pi et al., 2015). The control of QC activity and CSC maintenance by WOX5 also depends on cross-regulation and physical interactions with members of the PLETHORA (PLT) family of TFs (Burkart et al., 2022). As a result, *wox5* mutants display differentiated CSC but enhanced QC division to replenish shedding columella tiers, while WOX5 overexpression causes CC dedifferentiation (Bennett et al., 2014; Forzani et al., 2014; Sarkar et al., 2007).

Auxin also plays a critical role in the maintenance of distal stem cells through the formation of an instructive hormone gradient spanning from the QC throughout the CC (Ding and Friml, 2010; Dubreuil et al., 2018; Sabatini et al., 1999). The auxin gradient is generated and maintained through a combination of local hormone biosynthesis in the QC and polar transport within the distal tissues (Blilou et al., 2005; Stepanova et al., 2008). Interestingly, auxin biosynthesis in the QC is regulated by the action of WOX5 and in turn, high auxin levels promote CSC differentiation at least in part by downregulating WOX5 expression (Ding and Friml, 2010; Savina et al., 2020; Tian et al., 2014). Auxin-mediated WOX5 downregulation has been linked to the AUXIN RESPONSE FACTOR (ARF) 10 and ARF16 transcriptional regulators, which also play a role in restricting SCN and promoting CC differentiation (Ding and Friml, 2010; Wang et al., 2005). Double *arf10 arf16* mutants, or plants overexpressing *MIR160* - a microRNA which downregulates *ARF10* and *ARF16 -* display uncontrolled cell division and blocked cell differentiation in distal root tissues (Wang et al., 2005). Whether ARF10 and ARF16 are required for the auxin-mediated downregulation of WOX5 is still not clear, and it has been proposed that the two pathways act in parallel to regulate distal stem cells activity (Bennett et al., 2014).

Other than the auxin/WOX5 pathways, NAC transcription factors FEZ and SOMBRERO (SMB) are intrinsically required for correct developmental program of distal tissues (Bennett et al., 2014; Willemsen et al., 2008). Both *fez* and *smb* loss-of-function mutants display alterations in the number of columella and lateral root cap (LRC) cell layers from embryogenesis onward (Willemsen et al., 2008). In particular, *fez* mutants show a reduction in the number of columella and LRC layers due to lower frequency of CSC and epidermal Epi/LRC initials divisions, whereas *smb* mutants display additional CSC and LRC layers as a consequence of delayed differentiation (Willemsen et al., 2008). It has been proposed that FEZ and SMB operate in a self-regulatory loop to control CSC proliferation: FEZ is expressed in CSC where it induces ACD but is repressed by SMB in apical daughters to prevent further division and promote CC differentiation (Willemsen et al., 2008). Remarkably, this regulatory loop interacts with the WOX5 pathway, as it has been shown that WOX5 cell-autonomously represses FEZ expression in the QC to maintain quiescence, and non-cell-autonomously represses *SMB* in CSC to prevent differentiation (Bennett et al., 2014). Together, these studies clearly depict that distal stem cell activity does not rely on unified pathways regulating either proliferation or differentiation, but it rather depends on the convergence of multiple quantitative inputs, with presumably additional factors mediating regulatory interactions between known players (Bennett et al., 2014).

Here we identify members of the HD-Zip II transcription factor family as additional regulators of distal stem cell activity in roots. We show that developmental regulators HOMEOBOX ARABIDOPSIS THALIANA 3 (HAT3), ARABIDOPSIS THALIANA HOMEOBOX 4 (ATHB4), and ATHB2, known for their role in the control of apical embryo patterning and shoot architecture (Carabelli et al., 2021; Merelo et al., 2016; Turchi et al., 2013) redundantly regulate distal stem cell activity and columella organization. We provide evidence that HD-Zip II proteins regulate CSC activity by interfering locally with auxin signaling and at the same time by acting downstream of FEZ. Our work shows that HD-Zip II proteins control the balance between CSC proliferation and CC differentiation by bridging the gap between the FEZ/SMB and auxin pathways.

## RESULTS

### Simultaneous mutations of *HAT3*, *ATHB4*, and *ATHB2* affect RAM size and patterning

Previous work demonstrated that HAT3, ATHB4 and ATHB2 act redundantly in the regulation of shoot architecture (Carabelli et al., 2021; Merelo et al., 2016; Turchi et al., 2013), thus we wondered whether these TFs also play a role in root development. To this end, we characterized root phenotypes of single and double HD-Zip II mutant combinations. The root length and RAM size of *hat3-3 athb4-1* mutant seedlings were slightly although significantly reduced compared to the wild type (WT) (Figure S1). Similar defects were observed in different allelic combinations (*hat3-3 athb4-3, hat3-2 athb4-1)*, and fully rescued by introgressing an *HAT3::HAT3:GFP* construct in *hat3-3 athb4-1* (Figure S1). Single mutant analyses revealed that only *hat3-3* displays reduced root length and RAM size as *hat3 athb4*, suggesting that these phenotypes are not determined by the altered shoot development of the double mutant (Figure S1; Turchi et al., 2013). Closer inspection of *hat3-3 athb4-1* seedlings revealed defects in root tip organization. Irregular division patterns and abnormal QC and CC shapes were observed in more than 75% of the *hat3-3 athb4-1* root tips (Table S1) and at a similar level in the other allelic combinations. Root tip organization defects were only partially rescued by expressing *HAT3::HAT3:GFP* in *hat3-3 athb4-1* (Table S1). No significant alteration in single mutants’ root tip organization was observed (Table S1), indicating that HAT3 and ATHB4 function redundantly in this process. Next, we investigated whether lack of ATHB2 aggravates the *hat3 athb4* phenotype. Different allelic combinations of the *hat3 athb4 athb2* triple mutants displayed an increased severity of the root phenotypes observed in *hat3 athb4*. Reductions in root length and RAM size in *hat3 athb4 athb2* were already visible in 5-day-old seedlings, further reaching a dramatic decrease in 10-day-old plantlets (Figure 1, Table S1); defects in QC/CC patterning were observed at a higher frequency in *hat3 athb4 athb2* compared to double mutant combinations (Figure 1, Table S1). These phenotypes are caused by the concurrent lack of the three TFs since *athb2-1* loss-of-function mutants did not show differences in root length, RAM size and organization compared to the WT (Figure S1, Table S1). Also, lack of ATHB2 did not aggravate the phenotype of *hat3* mutants (Figure S1, Table S1). Only *athb4-1 athb2-3* displayed defects in RAM organization similar to those observed in *hat3 athb4*, although with reduced severity (Table S1). Interestingly, the introgression of *athb2-2* gain-of-function mutation or *ATHB2::ATHB2:GUS* construct in *hat3-3 athb4-1* were able to fully rescue the root length and RAM size without complementing QC/CC patterning defects (Figure S1; Table S1).

**Figure 1.**
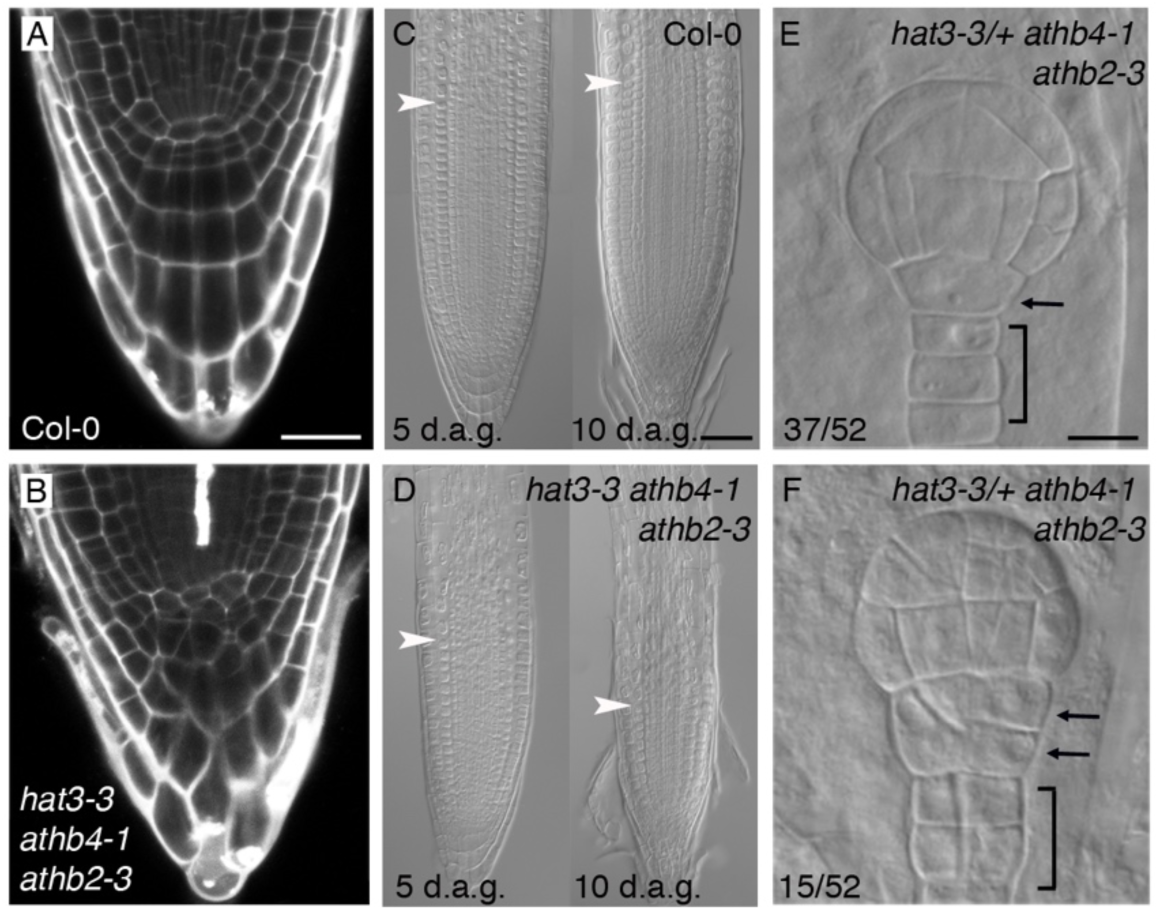
HD-Zip II TFs control root patterning. (A,B) Altered stem cell niche patterning in *hat3-3 athb4-1 athb2-3* (B) compared to Col-0 (A) visualized by Propidium Iodide (PI) staining. (C,D) Reduced meristem length in *hat3-3 athb4-1 athb2-3* (D) compared to Col-0 roots imaged at 5 and 10 days after germination (d.a.g). White arrowheads mark meristem boundaries. (E, F) Globular embryos from a segregating *hat3-3/+ athb4-1 athb2-3* genotype showing altered root pole in ∼25% of the population (F) compared to unaltered seedlings (E). Arrows and brackets mark the hypophysis and suspensor, respectively. Notice enhanced cell divisions in (F). Scale bars: A,B 5 μm; C,D 50 μm; E,F 20 μm

We next wondered whether the irregular organization of stem cells in *hd-zip II* mutants was caused by altered development of the embryo root pole. To this end, we analyzed early embryos from heterozygous *hat3-3 athb4-1/+* and *hat3-3/+ athb4-1 athb2-3* genotypes. We restricted the analysis to embryos up to the transition stage since the asymmetric division of the hypophysis that specifies the root pole occurs at this stage in the WT (Friml et al., 2003). In the progeny of *hat3-3 athb4-1/+*, about 25% of the embryos already displayed aberrant divisions of the hypophysis as expected by the segregation of a single recessive mutation (Figure S2). In the segregating progeny of *hat3-3/+ athb4-1 athb2-3* the phenotype was more severe as approximately one-quarter of the embryos displayed aberrant divisions of the hypophysis and of suspensor cells (Figure 1). Together, these data indicate that HAT3, ATHB4, and ATHB2 regulate root patterning and development from embryogenesis onward.

### HAT3, ATHB4, and ATHB2 are expressed in the RAM

To investigate HD-Zip II expression patterns in the RAM, we generated transgenic lines expressing fluorescent protein fusions for each TF. Because *HAT3::HAT3:GFP* was unable to fully rescue *hat3 athb4* root phenotypes (Figure S1; Table S1), we generated constructs containing the entire genomic *HAT3*, *ATHB4,* and *ATHB2* sequences, including the 3’ untranslated region (UTR), under the control of their specific promoters (respectively 4.7 Kbp, 4.6 Kbp and 5.4 Kbp upstream of ATG), in which the fluorescent protein marker was inserted with a poly-Alanine linker between the stop codon and the 3’ UTR (*HAT3::HAT3:YFP; ATHB4::ATHB4:GFP*; *ATHB2::ATHB2:GFP*; Figure S3A-C) Confocal analyses of HAT3:YFP, ATHB4:GFP and ATHB2:GFP expression patterns in the RAM showed that the three HD-Zip II proteins were predominantly expressed in the vascular tissue. ATHB2:GFP and HAT3:YFP were also detected in the pericycle (Figure 2A-G; Table S2). In distal tissues, the expression patterns of HD-Zip II TFs were largely overlapping, although with major differences in protein levels. Of the triplet, ATHB2:GFP displayed the highest expression, whereas HAT3:YFP and ATHB4:GFP levels were extremely low, often at the limit of confocal microscopy detection. Relevantly, the differences in the expression of HD-Zip II marker lines are coherent with tissue-specific transcript levels for each gene (Figure S3D). ATHB2:GFP expression was detected in CSC, the first two layers of CC and, at a minor frequency in the QC and Epi/LRC initials (Figure 2C,F,G; Table S2). ATHB4:GFP expression was similar to that of ATHB2:GFP except for the absence of signals in the QC and Epi/LRC initials (Figure 2B,E,G; Table S2). HAT3:YFP signals were observed in CSC and CC with an expression peak in the last columella tier. HAT3:YFP was also detected in the epidermis, LRC, and their initials (Figure 2A,D,G; Table S2). It is worth mentioning that HD-Zip II expression in distal RAM is highly dynamic, as we rarely observed homogeneous fluorescent signals in all the nuclei within these tissues, in particular in CSC. To better understand HD-Zip II dynamics in distal stem cells, we focused on ATHB2:GFP because of its relatively high expression. We observed that ATHB2:GFP is induced in CSC before the ACD and it is switched off in the resulting apical cell thereafter, while remaining expressed in the basal daughter which will later differentiate to CC (Figure S4).

**Figure 2.**
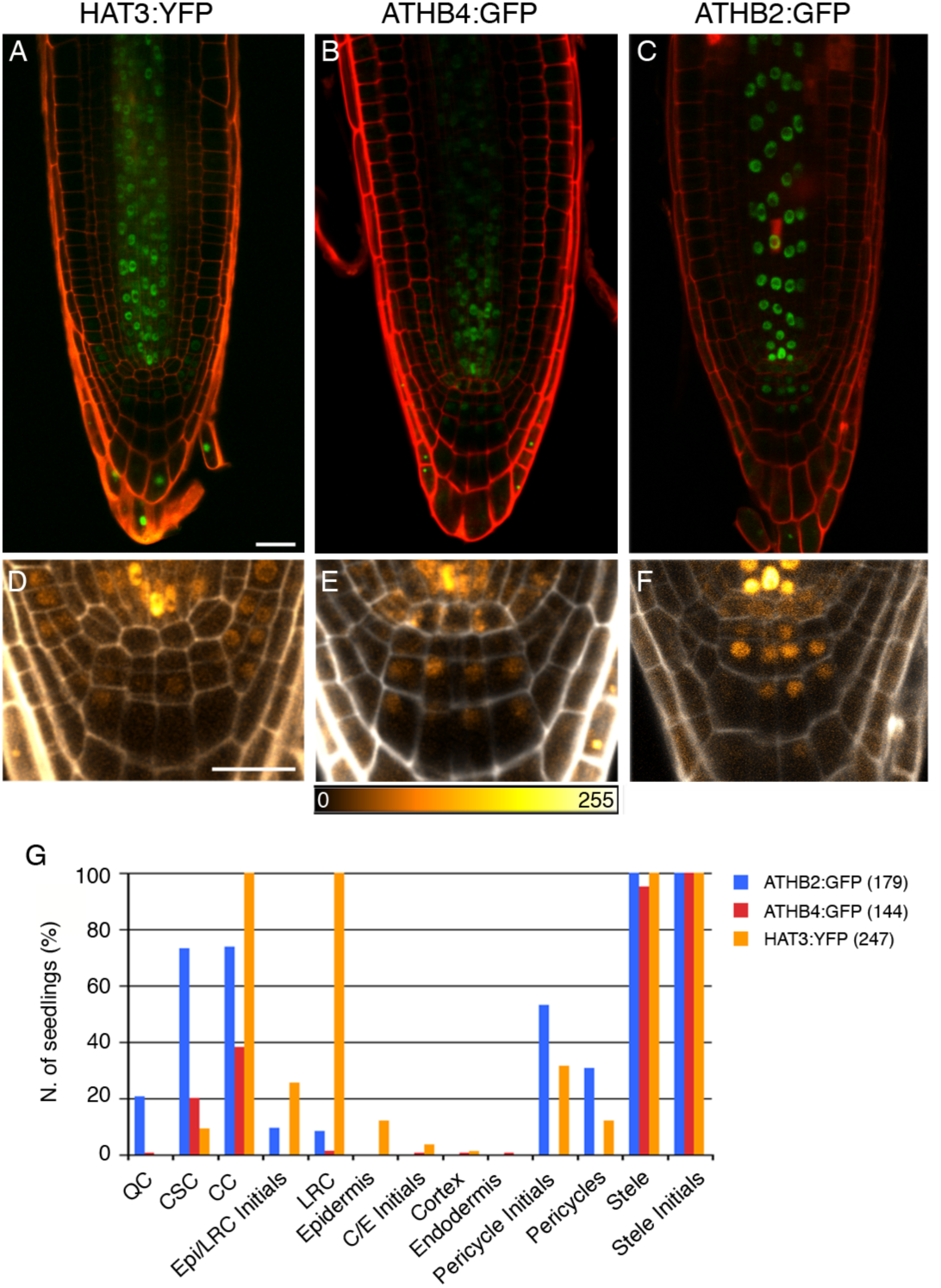
HD-Zip II are expressed in the root meristem. (A-C) Expression of HAT3:YFP (A), ATHB4:GFP (B) and ATHB2:GFP (C) proteins (green) in wild-type roots. (D-F) Details of HAT3:GFP (D), ATHB4:GFP (E) and ATHB2:GFP (F) expression in distal tissues of the same roots in (A-C), using the color scale indicated below. PI counterstaining, red (A-C), gray (D-F). Scale bars: 20 μm. (G) Occurrence of HD-Zip II reporter expression within meristematic tissues. The graph shows the percentage of roots displaying fluorescence in indicated cell types for each marker. Sample size is in brackets. C/E, Cortex/Endodermis.

Together, these data indicate that distal RAM phenotypes observed in multiple HD-Zip II mutants are likely caused by the lack of cell autonomous functions of these factors, regardless of their rather low expression levels. This is further supported by the fact that HAT3:GFP, which is unable to complement QC/CC patterning defects in *hd-zip II* mutants, is not detectable in distal root tissues (Figure S3F). The distal patterning defects, together with ATHB2:GFP expression dynamics in these tissues suggest that HD-Zip II function is related to distal stem cell divisions.

### HAT3, ATHB4, and ATHB2 regulate columella stem cell maintenance

To better understand HD-Zip II role in distal stem cells, we carried out Lugol’s starch staining to analyze the extent of CC differentiation in *hd-zip II* mutants. Depending on the phase of the cell cycle, wild-type columella contains one or two layers of CSC, the one farther from the QC is constituted by CSC-like cells. These are undifferentiated CSC daughter cells that will shortly expand and accumulate starch as part of their differentiation into CC (Bennett et al., 2014). Compared to the WT, *hat3 athb4* mutants display a reduced numbers of roots with 2 layers of CSC, an increased number of roots with one layer, and the appearance of roots with no CSC, as deduced by the appearance of starch granules in cells adjacent to the QC. This phenotype was completely rescued by introducing *HAT3::HAT3:YFP* in the double mutant background (Figure 3E). In *hat3-3 athb4-1 athb2-3* mutants, we observed the complete absence of RAMs with two layers of CSC and a significant increase in the frequency of those without CSC (Figure 3B,C,E). These phenotypes were not observed in *hat3-3 athb2-3* and *athb4-1 athb2-3* mutants, indicating that the presence of either HAT3 or ATHB4 is sufficient for CSC maintenance (Figure S5). Consistently, the expression of ATHB4:GFP in *hat3-3 athb4-1 athb2-3* restored the CSC phenotype to WT levels (Figure 3E). In contrast, the expression of ATHB2:GFP in *hat3-3 athb4-1 athb2-3* could only restore the phenotype to the levels of *hat3-3 athb4-1* (Figure 3E). Notably, it should be emphasized that the gain-of-function *athb2-2,* which expresses increased levels of *ATHB2* (Turchi et al., 2013), displays a higher percentage of roots with two layers of CSC compared to the WT (Figure 3A,D,E).

**Figure 3.**
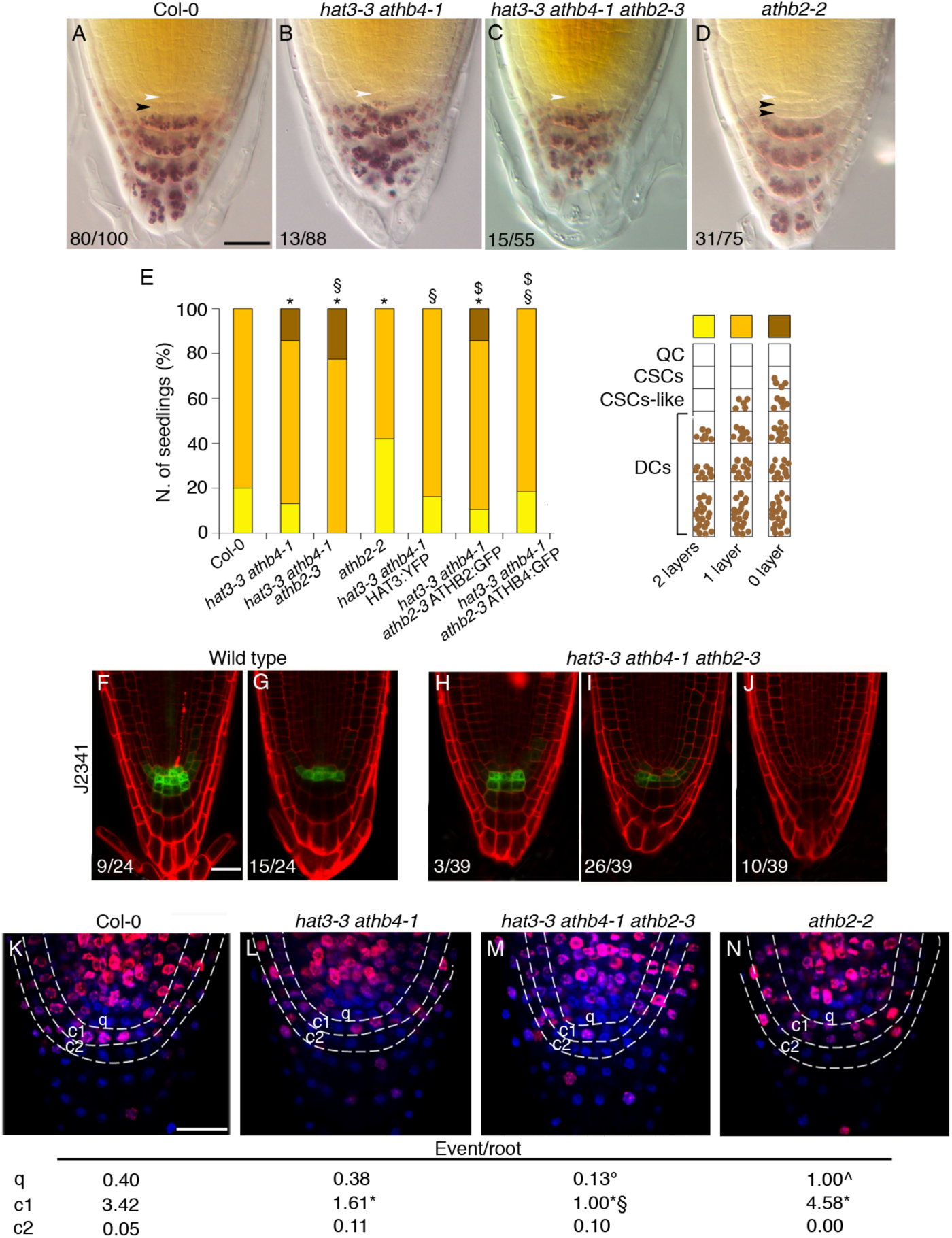
HD-Zip II regulate columella stem cells fate and identity. (A-D) CSCs differentiation is enhanced in *hat3-3 athb4-1* (B), *hat3-3 athb4-1 athb2-3* (C) and delayed in *athb2-2* (D) compared to Col-0 (A), as deduced by Lugol’s staining. White arrowheads, QC; black arrowheads, CSCs. (E) Quantification of CSC layers in the indicated genotypes, according to the scheme on the right. *p<0.01 *vs.* Col-0; §p<0.01 *vs. hat3-3 athb4-1*; $p<0.01 *vs. hat3-3 athb4-1 athb2-3*. (F-J) J2341 CSCs marker expression (green) in wild-type (F,G) and *hat3-3 athb4-1 athb2-3* (H,I,J) PI-stained roots (red). The ratio indicates the occurrence of the depicted expression pattern. (K-N) F-ara-EdU incorporation (magenta) in Col-0 (K), *hat3-3 athb4-1* (L), *hat3-3 athb4-1 athb2-3* (M), and *athb2-2* (N) roots counterstained with DAPI (Blue). *q*, *c1, c2* indicate QC, CSC, CC, respectively. The event/root index (sum of division and replication/number of roots) highlights differences in S-phase progression. °p<0.05 *vs*. Col-0 *q*; ^p<0.01 *vs*. Col-0 *q*; *p<0.01 *vs.* Col-0 *c1*; §p<0.01 *vs. hat3-3 athb4-1 c1*. Scale bars: A-D, 5 μm; F-N, 20 μm.

The enhanced CC differentiation observed in *hd-zip II* mutants (Figure 3A,B,E), is suggestive of a reduction in CSC proliferation. Thus, we used the thymidine analogue F-ara-EdU (Bennett et al., 2014; Hong et al., 2015) to study cell cycle progression and proliferation in the columella. We analyzed F-ara-EdU incorporation in WT, *hat3-3 athb4-1*, *hat3-3 athb4-1 athb2-3,* and *athb2-2* mutants, and quantified the number of DNA replication and cell division events at the QC (*q*), CSC (*c1*), and the first layer of CC (*c2*) positions (Figure 3K-N). Compared to the WT, loss- (*hat3-3 athb4-1*, *hat3-3 athb4-1 athb2-3*) and gain-of-function (*athb2-2*) mutants showed reduced and enhanced cell cycle progression at *c1*, respectively (Figure 3K-N, Table 1). In addition, reductions in cell division at *c1* were observed in *hat3-3 athb4-1* and, to a greater extent, in *hat3-3 athb4-1 athb2-3* compared to the WT. Noteworthy, *athb2-2* gain-of-function mutant displayed enhanced cell cycle progression and cell division also at *q* position, indicating that moderate increases in *ATHB2* expression are able to promote CSC and QC proliferation (Table 1). Together, Lugol’s and F-ara-EdU analyses show that HD-Zip II proteins play a role in CSC maintenance. Consistently, the expression of the CSC-specific marker J2341 was severely compromised in *hat3-3 athb4-1 athb2-3* indicating the loss of stem cell identity in absence of HD-Zip II (Figure 3F-J).

**Table 1.** Quantification of F-ara-EdU incorporation in the genotypes used in this study.

|  | Position | Replication | Division | Event/root |
| --- | --- | --- | --- | --- |
| Col-0<br>n=35 | *q* | 14 | 0 | 0.40 |
|  | *c1* | 56 | 64 | 3.42 |
|  | *c2* | 2 | 0 | 0.05 |
| *hat3-3 athb4-1*<br>n=36 | *q* | 14 | 0 | 0.38 |
|  | *c1* | 41 | 17 | 1.61* |
|  | *c2* | 4 | 0 | 0.11 |
| *hat3-3 athb4-1 athb2-3*<br>n=30 | *q* | 4 | 0 | 0.13° |
|  | *c1* | 22 | 8 | 1.00* |
|  | *c2* | 3 | 0 | 0.10 |
| *athb2-2*<br>n=26 | *q* | 18 | 8 | 1.00^ |
|  | *c1* | 59 | 60 | 4.58* |
|  | *c2* | 0 | 0 | 0.00 |
| *wox5-1*<br>n=32 | *q* | 35 | 14 | 1.53# |
|  | *c1* | 28 | 4 | 1.00* |
|  | *c2* | 3 | 0 | 0.09 |
| *hat3-3 athb4-1 athb2-3*<br>*wox5-1*<br>n=26 | *q* | 21 | 12 | 1.26# |
|  | *c1* | 12 | 0 | 0.46* ∫ ^ |
|  | *c2* | 2 | 0 | 0.07 |
| *fez-2*<br>n=28 | *q* | 15 | 2 | 0.60 |
|  | *c1* | 30 | 9 | 1.39* |
|  | *c2* | 6 | 0 | 0.21 |
| *hat3-3 athb4-1 athb2-3*<br>*fez-2*<br>n=31 | *q* | 9 | 0 | 0.29 |
|  | *c1* | 30 | 2 | 1.03* |
|  | *c2* | 4 | 0 | 0.12 |
| *smb-3*<br>n=27 | *q* | 15 | 0 | 0.55 |
|  | *c1* | 58 | 22 | 2.96 |
|  | *c2* | 35 | 9 | 1.62« |
| *hat3-3 athb4-1 athb2-3*<br>*smb-3*<br>n=31 | *q* | 6 | 0 | 0.19° |
|  | *c1* | 27 | 6 | 1.06* |
|  | *c2* | 5 | 1 | 0.19∞ |

The number of DNA replication and cell division events was quantified by counting F-ara-EdU positive nuclei at indicated positions. The final column shows sum of replications and divisions, divided by the number of roots. °p<0.05 vs. Col-0 *q*; #p<0.01 vs. Col-0 *q*; \*p<0.01 vs. Col-0 *c1*; ∫ p<0.05 vs. *hat3-3 athb4-1 athb2-3 c1*; ^p<0.05 vs. *wox5-1 c1*; §p<0.01 vs. *hat3-3 athb4-1 c1*; «p<0.01 vs. Col-0 *c2*; ∞p<0.01 vs. *smb-3*.

Overall, these data demonstrate that HD-Zip II TFs maintain the identity and function of CSC by sustaining their proliferative status. The genetic analyses demonstrate that the presence of either HAT3 or ATHB4, but not of ATHB2, is necessary for CSC maintenance. However, ATHB2 function positively correlates with stem cell proliferation, as higher *ATHB2* levels in the gain-of-function mutant increase QC e CSC proliferation.

### HAT3, ATHB4 and ATHB2 interfere with auxin response to regulate CSC activity

Auxin regulates the formation of the columella during the embryogenesis, as well as its maintenance and differentiation in post-embryonic roots (Ding and Friml, 2010; Dubreuil et al., 2018; Friml et al., 2003; Weijers et al., 2006). We thus wondered whether distal RAM defects observed in *hd-zip II* mutants were due to altered auxin distribution or response. We analyzed the expression pattern of DR5rev::GFP auxin response marker in relation with aberrant hypophysis division in *hat3-3/+ athb4-1* segregating embryos. In globular embryos displaying altered root pole, DR5rev::GFP expression was markedly reduced in hypophysis and increased in suspensor cells, as opposed to WT-like embryos which only displayed DR5 signals in the hypophysis (Figure S6). These results suggest that altered root pole development in *hat3 athb4* is caused by a defective establishment of auxin maxima. However, in *hat3 athb4* mature embryos (Turchi et al., 2013), and in post-embryonic roots DR5rev::GFP expression does not differ substantially from the WT (Figure S6).

Next, we analyzed DR5rev::GFP post-embryonic expression pattern in *hat3 athb4 athb2* mutant roots. Despite the severe CSC phenotypes, *hat3-3 athb4-1 athb2-3* did not display at a first glance major differences in DR5 expression compared to the WT (Figure 4A,B). However, upon quantitative analyses of DR5 expression, we could highlight differences between the WT and the triple mutant. In particular, in *hat3 athb4 athb2* the peak of DR5 expression shifted from the QC to the CSC (Figure 4A,B). Because the expression pattern of the auxin biosynthesis reporter TAA1::GFP:TAA1 did not appreciably change in *hat3 athb4 athb2* compared to the WT (Figure S7), we wondered whether the shift of DR5 expression could be attributable to altered auxin transport. To this end, we analyzed the expression of PIN3, PIN4 and PIN7 auxin efflux carriers in *hat3-3 athb4-1 athb2-3*. We found that the expression of PIN3:GFP and PIN4:GFP, but not that of PIN7:GFP, was enhanced in the triple mutant compared to the WT (Figure 4C-I). The increase in PIN3 and PIN4 levels was particularly evident in distal tissues where they play a key role in the maintenance of auxin maximum and columella differentiation (Blilou et al., 2005; Ding and Friml, 2010). To test whether altered PINs expression contributes to *hd-zip II* distal stem cells defects, we analyzed CSC phenotypes in multiple *hd-zip II pin* mutant combinations. In contrast to *pin3-4, pin4-3, pin7-2* mutants, showing higher frequencies of RAMs with two CSC layers compared to the WT (Blilou et al., 2005; Ding and Friml, 2010; Friml et al., 2002b), quadruple mutant combinations *hat3-3 athb4-1 athb2-3 pin3-4*, *hat3-3 athb4-1 athb2-3 pin4-3*, and *hat3-3 athb4-1 athb2-3 pin7-2* displayed enhanced CSC differentiation like the triple *hd-zip II* (Figure 4J). Thus, increased expression of PINs in CC is unlikely the cause of enhanced CSC differentiation in *hat3 athb4 athb2*.

**Figure 4.**
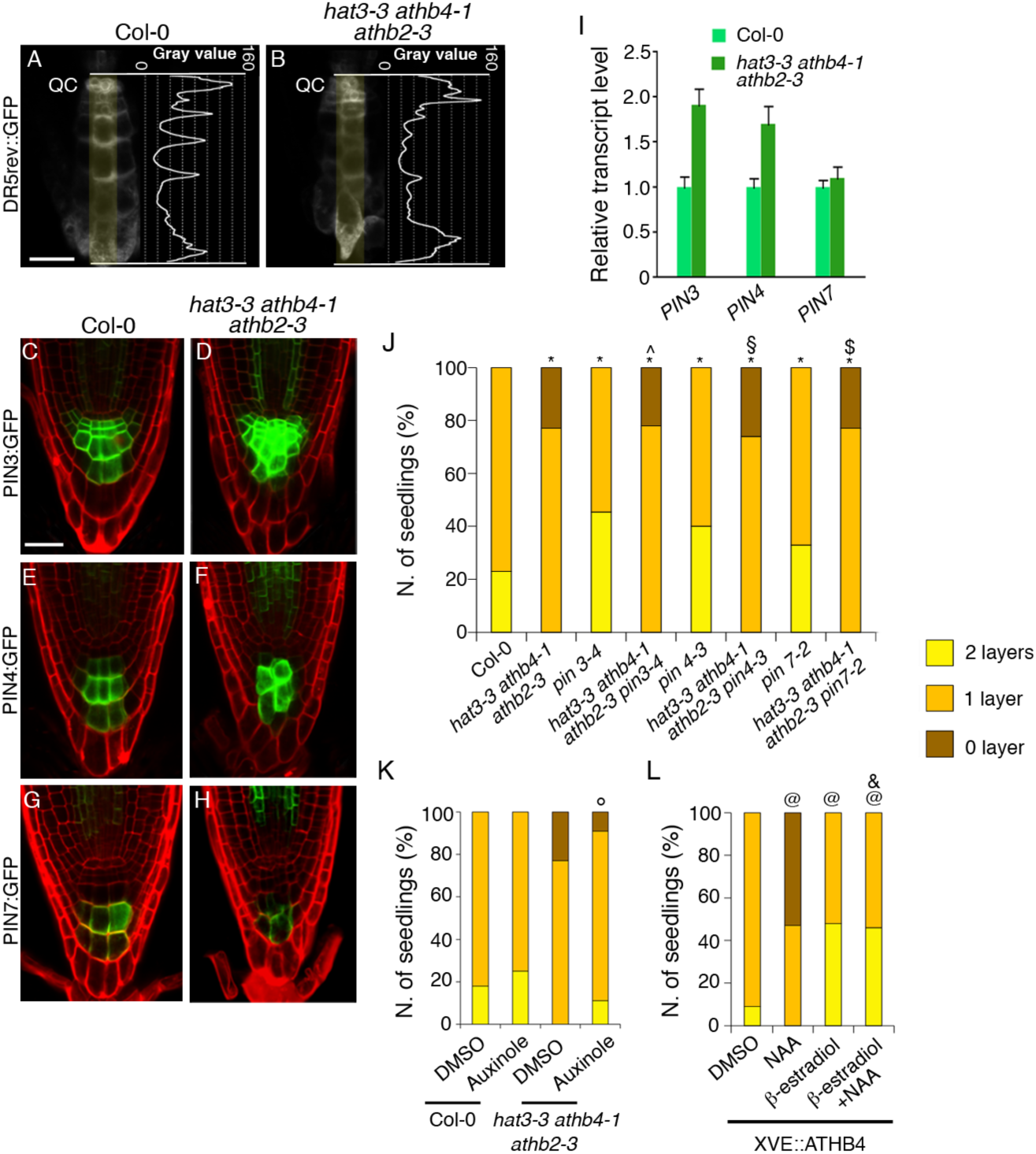
HD-Zip II affect auxin transport and signaling in the RAM. (A,B) Altered *DR5rev::GFP* expression in *hat3-3 athb4-1 athb2-3* (B) compared with Col-0 (A). Signal intensity profiles measured along the highlighted regions are reported on the right. (C-I) Altered PIN expression in *hd-zip II* mutants. (C-I) PIN3:GFP (C,D), PIN4:GFP (E,F) and PIN7:GFP (G,H) expression in Col-0 (C,E,G) and *hat3-3 athb4-1 athb2-3* (D,F,H) PI-Stained roots. Scale bars: 20μm. (I) Relative expression levels of *PIN3*, *PIN4* and *PIN7* in Col-0 and *hat3-3 athb4-1 athb2-3* roots determined by qRT-PCR. Error bars, standard deviation (s.d.). (J) Quantification of CSC layers in multiple *hd-zip II pin* mutant combinations. *p<0.01 *vs.* Col-0; ^p<0.01 *vs*. *pin3-4*; §p<0.01 *vs*. *pin4-3*; $p<0.01 *vs*. *pin7-2*. (K) Inhibition of auxin signaling mitigates *hat3 athb4 athb2* CSCs phenotype. Quantification of CSC layers in Col-0 or *hat3-3 athb4-1 athb2-3* treated with DMSO or 5μM auxinole. °p<0.01*vs. hat3-3 athb4-1 athb2-3* DMSO. (L) Induction of ATHB4 inhibits auxin-mediated CSC differentiation. Quantification of CSC layers in in *XVE>ATHB4* treated as indicated. @p<0.01 *vs.* DMSO; &p<0.01 *vs.* NAA.

We next tested whether alterations in auxin signaling mediated by TRANSPORT INHIBITOR RESISTANT 1/AUXIN BINDING F-BOX (TIR1/ABF) receptors could explain *hd-zip II* mutant phenotypes. We analyzed the effect of auxinole, a potent auxin antagonist of TIR1/AFBs receptors (Hayashi et al., 2008; Hayashi et al., 2012), on WT and *hat3 athb4 athb2* roots. In the WT, auxinole treatments increased the frequency of RAMs with 2 layers of CSC compared to untreated controls, in agreement with an expected inhibition of auxin-mediated CC differentiation. In *hat3-3 athb4-1 athb2-3*, auxinole restored the occurrence of RAMs with 2 layers of CSC normally absent in this background (Figure 4K), suggesting that HD-Zip II TFs might regulate CSC activity by counteracting auxin signaling. Thus, we expressed ATHB4 under the control of a β-estradiol inducible system (XVE>ATHB4) to test whether high levels of an HD-Zip II TF counteract the auxin-mediated CSC differentiation. Consistently with the phenotype of the gain-of-function *athb2-2*, high levels of ATHB4 enhanced CSC proliferation (Figure 4L; Figure S8) and cell cycle progression at *c1* in β-estradiol-treated vs. untreated plants (Figure S8). Remarkably, the application of NAA did not enhance CSC differentiation upon induction of ATHB4, as observed in uninduced controls (Figure 4L).

Together, these data indicate that HD-Zip II regulate CSC fate by affecting the cellular readout of auxin signaling.

### HD-Zip II interfere with the ARF10/ARF16 pathway, but not with WOX5, to regulate CSC activity downstream of auxin

Auxin-mediated CSC differentiation has been linked to the action of WOX5 (Bennett et al., 2014; Ding and Friml, 2010; Savina et al., 2020). To understand whether WOX5 contributes to HD-Zip II functions in distal stem cells, we analyzed WOX5::GFP expression in *hat3 athb4 athb2* and found that it was substantially unchanged compared to the WT (Figure 5A,B). We thus wondered whether HD-Zip II, which act as transcriptional repressors (Steindler et al., 1999), could be involved in the auxin-mediated downregulation of WOX5 expression. Interestingly, we observed that ATHB2:GFP, is significantly upregulated by NAA treatments in the RAM (Figure 5 J,K). In distal tissues, NAA increased the frequency of QC and CSC cells expressing ATHB2:GFP at a given time (Figure 5K; Figure S9), suggesting that auxin stabilizes ATHB2 expression dynamics in distal tissues. However, WOX5::GFP transcriptional response to NAA treatment was not different between the WT and the triple mutant (Figure 5C,D) indicating that the auxin-mediated WOX5 downregulation is independent of HD-Zip II. Consistently, Lugol’s and F-ara-Edu analyses demonstrated additivity of the *hat3 athb4 athb2 wox5* mutant phenotypes, indicating that HD-Zip II and WOX5 regulate CSC activity through parallel pathways (Figure 5E-I).

**Figure 5.**
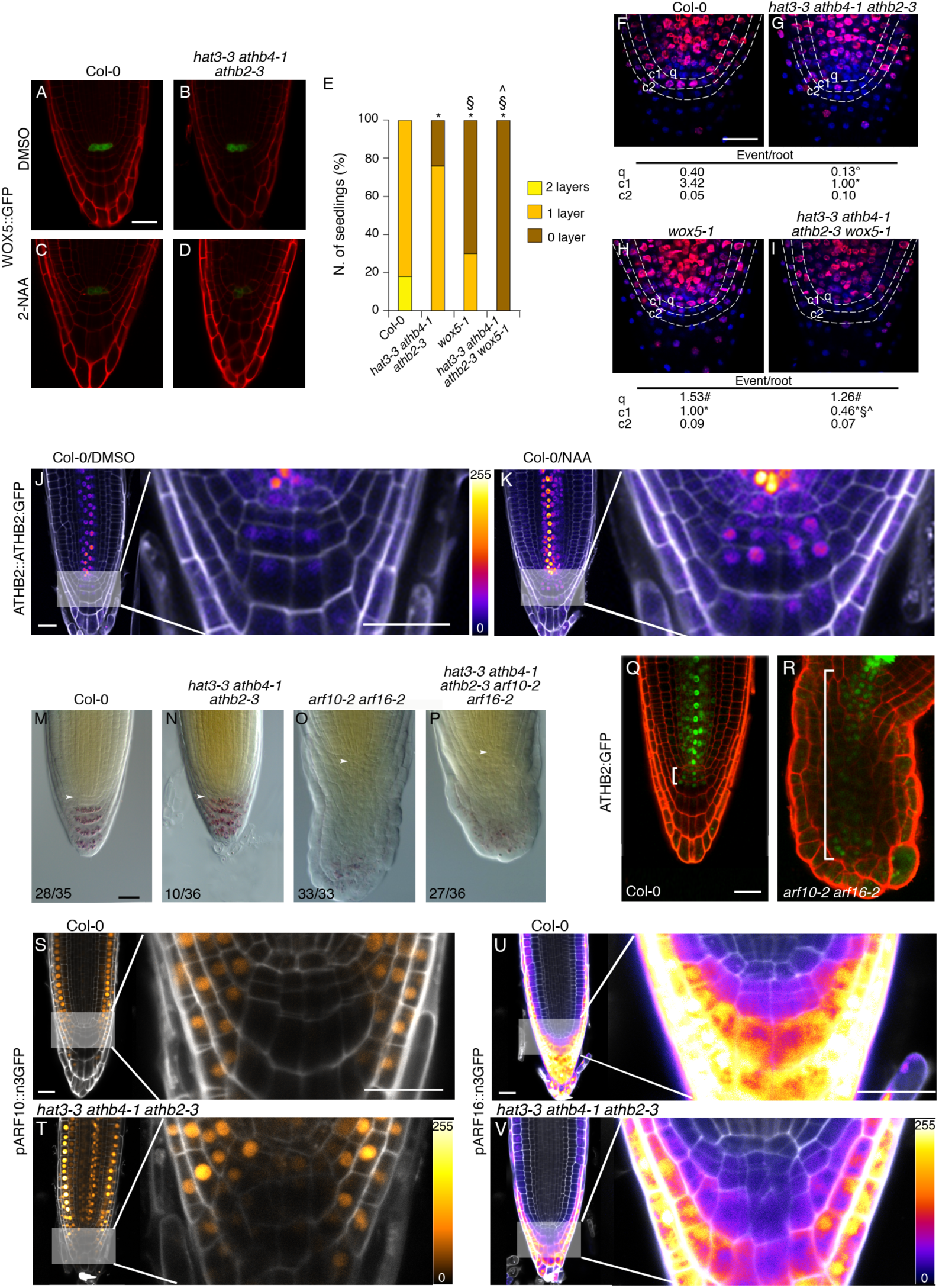
Genetic interactions between HD-Zip II and auxin-related CSC regulators WOX5 and ARF10/ARF16. (A-I) HD-Zip II act independently of WOX5. WOX5::GFP expression in Col-0 (A,C), and *hat3-3 athb4-1 athb2-3* (B,D) roots, untreated (A,B), or treated with NAA (C,D). (E) Quantification of CSC layers in *hat3-3 athb4-1 athb2-3 wox5-1* showing phenotype additivity compared to parental genotypes. *p<0.01 *vs*. Col-0; §p<0.01 *vs. hat3-3 athb4-1 athb2-3; ^*p<0.01 *vs. wox5-1*. (F-I) F-ara-EdU incorporation (magenta) in Col-0 (F), *hat3-3 athb4-1 athb2-3* (G), *wox5-1* (H), *hat3-3 athb4-1 athb2-3 wox5-1* (I) roots counterstained with DAPI (Blue). The event/root index highlights differences in S-phase progression. °p<0.05 *vs*. Col-0 *q*; #p<0.01 *vs*. Col-0 *q*; *p<0.01 *vs*. Col-0 *c1*; §p<0.05 *vs*. *hat3-3 athb4-1 athb2-3 c1*; ^p<0.05 *vs*. *wox5-1 c1*. (J, K) Expression ATHB2:GFP in roots treated for 24h with DMSO (J) or NAA (K). Magnifications show ATHB2:GFP increase in SCN upon NAA treatment. GFP is displayed using the color scale in inset, PI in gray. (M-P) Genetic interactions between *hd-zip II* and *arf* mutants. Lugol-stained roots of Col-0 (M), *hat3-3 athb4-1 athb2-3* (N), *arf10-2 arf16-2* (O) and *hat3-3 athb4-1 athb2-3 arf10-2 arf16-2* (P). White arrowheads indicate QC. (Q, R) *ATHB2:GFP* expression (green) in Col-0 (Q) and *arf10-2 arf16-2* (R) PI-stained roots (red). Brackets indicate ATHB2 expression domains. (S-V) Expression of pARF10::n3GFP (S-T) and pARF16::n3GFP (U,V) in Col-0 (S,U) and *hat3-3 athb4-1 athb2-3* (T,V) roots. Magnifications show changes in ARFs expression domains in the SCN. GFP is displayed using the color scale in insets, PI is in gray. Scale bars: A-D, F-K, Q-V 20 μm; M-P, 5 μm.

To further investigate the link between auxin and HD-Zip II, we focused on ARF10 and ARF16 whose double loss-of-function mutant displays uncontrolled cell division and impaired cell differentiation in the distal RAM (Wang et al., 2005). To test the genetic relationship between HD-Zips II and ARF TFs, we generated and analyzed the *hat3-3 athb4-1 athb2-3 arf10-2 arf16-2* quintuple mutant. The phenotype of *hat3-3 athb4-1 athb2-3 arf10-2 arf16-2* showed mitigation of the uncontrolled cell proliferation with fewer layers of undifferenyiated cells with CSC morphology compared to the parental *arf10 arf16* (Figure 5M-P), suggesting that HD-Zip II contribute to the increased CSC proliferation of *arf10 arf16*. Consistently, we found that ATHB2:GFP is expressed in an enlarged domain of *arf10 arf16* distal RAM compared to the WT (Figure 5Q,R). Also, we found that pARF10::n3GFP expression domain, which is restricted to the LRC/Epidermis initials in distal tissues of the WT, expands toward CSC and CC in *hat3-3 athb4-1 athb2-3* (Figure 5S,T). Conversely, the expression of pARF16::n3GFP is specifically downregulated in the CSC/CC of the *hat3 athb4 athb2* mutant compared to the WT (Figure 5U,V). Together these data suggest an interdependency of HD-Zip II and ARFs in controlling the balance between proliferation and differentiation in distal stem cells.

### HD-Zip II act downstream of FEZ/SMB loop to control CSC proliferation

Since the genetic interaction with ARFs cannot fully explain the phenotypes observed in *hat3 athb4 athb2*, we next tried to determine whether HD-Zip II connect with FEZ and SMB, key players of stem cell fate and patterning in the root cap (Willemsen et al., 2008). To gain insights on the interactions among HD-Zip II proteins and FEZ/SMB pathway, we generated the quadruple mutants *hat3-3 athb4-1 athb2-3 fez-2* and *hat3-3 athb4-1 athb2-3 smb-3.* Lugol’s analysis showed that the *hat3-3 athb4-1 athb2-3 fez-2* quadruple mutant displays the same phenotype as *fez-2* (Figure 6A). F-ara-EdU experiments showed that cell cycle progression at *c1* is slightly lower, although not in a significant manner, in *hat3-3 athb4-1 athb2-3* than in *fez-2* with respect to WT. Also, the proliferative activity of *hat3-3 athb4-1 athb2-3 fez-2* resembles that of *hat3-3 athb4-1 athb2-3* (Figure 6 B,C,D,E; Table 2), suggesting that HD-Zip II and FEZ act in the same pathway. To further investigate the relationship between HD-Zips II and FEZ, we analyzed the expression of FEZ:GFP in *hat3-3 athb4-1 athb2-3* and found no significant difference between the triple mutant and the WT (Figure 6H,I). On the other hand, ATHB2:GFP was not expressed in root cap tissues of the *fez-2* mutant (Figure 6J,K). Importantly, ATHB2:GFP expression could not be restored in distal tissues even in presence of NAA (Figure 6M,N), indicating that FEZ is required for the auxin-mediated induction of ATHB2. Based on these results, we hypothesized that the strong reduction in cell cycle progression at *c1* in *fez* mutant might be caused by the loss of *HD-Zip II* expression. To verify this hypothesis, we introduced in the *fez-2* background the *35S::HAT3:GR* construct expressing a dexamethasone (DEX)-inducible version of HAT3, which like ATHB2 is not present in *fez-2* CSC (Figure S10). Upon DEX induction, HAT3:GR was able to fully complement *hat3-3 athb4-1 athb2-3* CSC phenotypes, confirming the functionality of the chimeric protein (Figure 6O-Q). Remarkably, HAT3:GR was able to largely recover the CSC phenotype of *fez-2* upon DEX induction (Figure 6O-Q), indicating that FEZ activity on CSC proliferation largely depends on HD-Zip II TFs.

**Figure 6.**
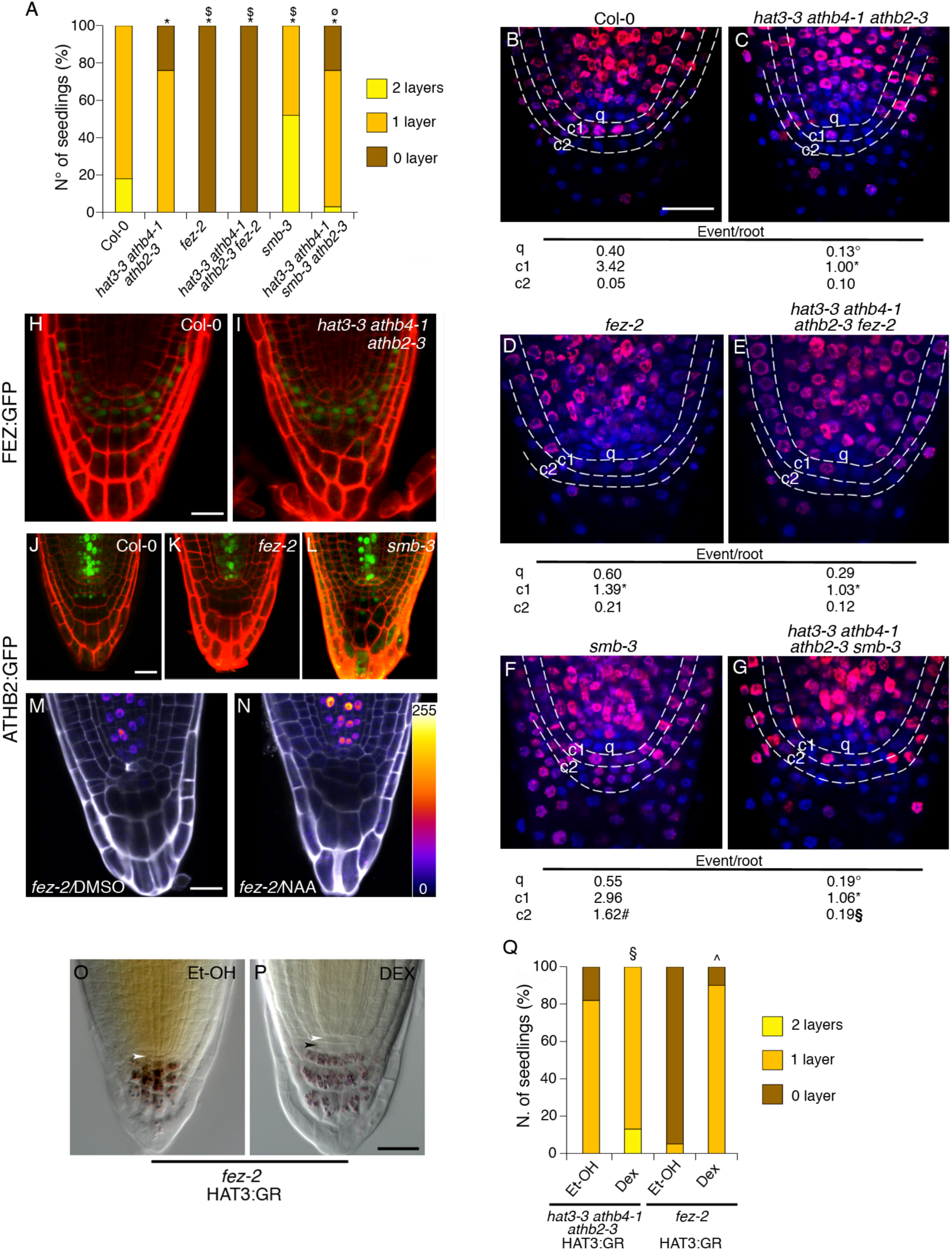
HD-Zips II TFs act downstream the FEZ/SMB pathway to regulate CSCs fate. (A) Quantification of CSC layers in *hd-zip II*, *fez* and *smb* mutant combinations. *p<0.01 *vs*. Col-0; $p<0.01 *vs. hat3-3 athb4-1 athb2-3*; ^ø^p<0.01 *vs. smb-3*. (B-G) F-ara-EdU incorporation (magenta) in Col-0 (B), *hat3-3 athb4-1 athb2-3* (C) *fez-2* (D), *hat3-3 athb4-1 athb2-3 fez-2* (E), *smb-3* (F) and *ha3-3 athb4-1 athb2-3 smb-3* (G) roots counterstained with DAPI (Blue). The event/root index highlights differences in S-phase progression. °p<0.05 *vs*. Col-0 *q*; *p<0.01 *vs*. Col-0 *c1*; #p<0.01 *vs*. Col-0 *c2*; §p<0.01 *vs*. *smb-3 c2*. (H, I) No difference in FEZ:GFP expression between Col-0 (H), and *hat3-3 athb4-1 athb2-3* (I). (J-L) Altered expression of ATHB2:GFP in *fez-2* (K) and *smb-3* (L) compared with Col-0 (J). (M,N) ATHB2:GFP expression in *fez-2* roots treated with DMSO (M) or NAA for 24h (N) according to the color scale in the inset, PI in gray. (O-P) *fez-2* HAT3:GR roots treated with EtOH (O) or dexamethasone (DEX) for 24h (P). White arrowheads, QC; black arrowheads, CSCs. (Q) Quantification of CSC layers in *hat3-3 athb4-1 athb2-3* HAT3:GR and *fez-2* HAT3:GR, treated with EtOH or DEX for 24h. *p<0.01 *vs*. *hat3-3 athb4-1 athb2-3* HAT3:GR/Et-OH; ^p<0.01 *vs*. *fez-2* HAT3:GR/Et-OH. Scale bars: B-N 20 μm; O-P 5 μm.

We next analyzed the interactions among HD-Zip II and SMB. Lugol’s analysis showed that CSC differentiation is delayed in *smb-3* mutant (Figure 6A). F-ara-Edu analysis showed that the number of mitotic events at *c2* is significantly increased in *smb-3* compared to the WT, implying that in absence of SMB CSC daughters maintain stem cell fate and undergo an extra round of division (Figure 6B,F; Table 2). In agreement with this, we found that in *smb-3* mutants ATHB2:GFP was predominantly expressed in *c2-c3* layers, following the proliferative pattern of distal cells (Figure 6J,L). Interestingly, enhanced proliferation of *smb-3* was recovered by the lack of HD-Zip II, as quadruple *hat3-3 athb4-1 athb2-3 smb-3* mutants display essentially the same phenotype of *hat3-3 athb4-1 athb2-3* (Figure 6B,C,F,G; Table 2). These results are in line with the idea that SMB drives CSC differentiation by excluding FEZ from the *c2* region (Willemsen et al., 2008), and with our finding that FEZ promotes CSC proliferation through the activity of HD-Zip II proteins. Together, the combination of genetic and expression data strongly suggests that HD-Zip II proteins act as effectors of FEZ/SMB loop in controlling CSC proliferation.

## Discussion

The maintenance of the SCN in the Arabidopsis root tip is precisely regulated by multiple, independent pathways that impinge quantitatively on the processes governing the switch between proliferation and differentiation (Bennett et al., 2014). Here, we provide evidence that HAT3, ATHB4, and ATHB2, originally identified as regulators of plant responses to canopy shade (reviewed in Ruberti et al., 2012; Sessa et al., 2018), are involved in the patterning of distal root tissues by controlling root pole formative divisions during embryogenesis, and CSC proliferation/differentiation balance in post-embryonic development. Regarding the latter aspect, our work shows that the three HD-Zip II play a role in promoting distal stem cell proliferation as confirmed by the enhanced CSC differentiation in double and triple mutants. Although HAT3, ATHB4, and ATHB2 seem not completely redundant in the process increased expression of either one consistently promotes CSC proliferation (Figure 3, 4 and 6; Table 1), indicating their direct role in the regulation of stem cell maintenance. HD-Zip II TFs do not function as another independent pathway, they rather take part in the regulatory circuitry underlying distal stem cell maintenance by connecting the auxin and the FEZ/SMB pathways.

The FEZ/SMB pathway represents a core mechanism setting the pace of CSC division/differentiation decision-making process and relies on a precise feedback loop between the two NAC TFs across adjoining root cap cell layers, with SMB promoting differentiation by repressing FEZ-mediated proliferative activity (Bennett et al., 2014; Willemsen et al., 2008). Interestingly, FEZ activity promotes CSC proliferation although the protein itself is not strictly required for cell division (Bennett et al., 2014). In this framework, our results suggest that the HD-Zip II act as effectors of FEZ in the regulation of CSC proliferation. Several lines of evidence support this hypothesis: HD-Zip II are epistatic to FEZ; FEZ is required for the expression of ATHB2; and inducing HAT3:GR expression restores CSC proliferation in *fez* mutants. It is worth mentioning that the cycling expression of ATHB2 in CSC and daughter cells is highly reminiscent of that described for FEZ (Willemsen et al., 2008), and the expanded expression domains of ATHB2 in *smb* mutants, where FEZ is not repressed, further suggest that FEZ could be directly involved in the regulation of HD-Zip II in distal tissues. Relevantly, FEZ activity has been shown to be sufficient to redirect the plane of cell division, even in tissues in which FEZ is not normally expressed, or during tissue regeneration processes (Marhava et al., 2019; Willemsen et al., 2008). Similarly, ectopic expression of ATHB4 and HAT3 promotes the formation of periclinal division in epidermis, cortex, endodermis and pericycle cells (Figure S8), suggesting that the interplay of HD-Zip II and FEZ could be active even outside the CSC and, most importantly, it could be involved in the regulation of cell division planes in the root pole during embryogenesis.

Our results further indicate that HD-Zip II TFs regulate distal stem cells proliferation by counteracting the auxin-induced CSC differentiation. Two main lines of evidence support this view: the enhanced differentiation of CSC in the triple *hd-zip II* mutant can be dampened by inhibiting auxin signaling; overexpressing ATHB4 blocks the effect of auxin on CSC differentiation. These data clearly indicate that HD-ZIP II TFs play a role in tuning down the cellular readout of auxin signaling in the root cap. Relevantly, the expression patterns of ARF10 and ARF16 are altered in the triple *hd-zip II* mutant, the first being upregulated, and the other downregulated in the mutant columella. ARF10 and ARF16 are auxin-inducible, class C (i.e. negative) ARFs that are expressed in partially overlapping domains and redundantly function to repress cell proliferation in the distal RAM (Dai et al., 2021; Mutte et al., 2018; Rademacher et al., 2011; Wang et al., 2005). ARF10 and ARF16 were proposed to repress CSC proliferation in response to auxin via WOX5 non-cell-autonomous function (Ding and Friml, 2010). Our genetic and molecular data show that HD-Zip II are unlikely to interfere with WOX5-mediated regulation of CSC cell fate. Instead, our data suggest that the HD-Zip II TFs, by controlling the expression domains of ARF10 and ARF16, modify the local perception of auxin levels, thus modulating the balance between proliferation and differentiation. It remains to be established how the opposite regulation of ARF10 and ARF16 by HD-Zip II TFs is achieved. A simple possibility is that HD-Zip II only target ARF10, which in turn controls ARF16 expression, in line with the current view of auxin responses being regulated by networks of transcriptional repressors (Truskina et al., 2021). It must be pointed out that the activity of ARF10 and ARF16 in the CSC has been demonstrated to be regulated by IAA33 (Lv et al., 2020). IAA33, like other non-canonical AUX/IAA proteins, is stabilized by auxin and regulates the transcriptional activity of ARFs in competition with canonical, auxin-degradable AUX/IAAs (Cao et al., 2019; Lv et al., 2020). Thus, it is possible that additional AUX/IAA-ARF pairs downstream of TIR1/ABFs (Lv et al., 2020; Wang et al., 2023), may be involved in the HD-Zip II-mediated regulation of auxin perception, as further suggested by the partial rescue of *hd-zip II* mutants phenotype by auxinole. Interestingly, IAA33 was identified as a putative interactor of ATHB2 in a large-scale yeast two-hybrid interactome screen (Wanamaker et al., 2017), opening the possibility for a more complex regulatory mechanism linking HD-Zip II activity to ARF-mediated auxin responses. It must be also considered that HD-Zip II TFs may interfere with auxin signaling indirectly, by altering local hormone fluxes via the transcriptional control on *PINs* (this work; Yuan et al., 2021).

Interestingly, the expression of ATHB2 is regulated by auxin, similarly to what observed for other members of the γ subfamily of HD-Zip II (He et al., 2020; Sawa et al., 2002). Auxin induces ATHB2 and stabilizes its expression dynamics in distal tissues. The auxin-mediated regulation of ATHB2 expression does not occur in *fez* mutants (Figure 6), suggesting that ATHB2 counteracts IAA-induced differentiation by promoting stem cell proliferation downstream of FEZ. Indeed, ATHB2 is also induced by auxin in the QC, where it stimulates divisions, as inferred by the gain-of-function mutant. The QC is known to divide to replenish differentiating CSC, acting as a reservoir of stem cells in conditions of extreme stress/damage (Cruz-Ramírez et al., 2013; Heyman et al., 2013). Thus, ATHB2 upregulation may represent a feedback response to prevent exaggerated CSC differentiation in the presence of excess auxin. Taken together, our data show that HD-Zip II TFs regulate distal stem cell fate by integrating opposing inputs to fine tune the balance between proliferation and differentiation (Figure 7). This integration may be of primary importance to synchronize root cap growth with the environment. Auxin acts as a messenger that regulates root growth in response to different environmental conditions, including light, temperature, drought, and nutrient deprivation (Ai et al., 2023; Hong et al., 2017; Liu and von Wirén, 2022; Sassi et al., 2012). In this framework, HD-Zip II TFs may act as a hub that integrates developmental cues (i.e., FEZ/SMB loop) with auxin-transmitted environmental cues to regulate distal stem cell behavior and, as a result, the dynamics of root cap growth.

**Figure 7.**
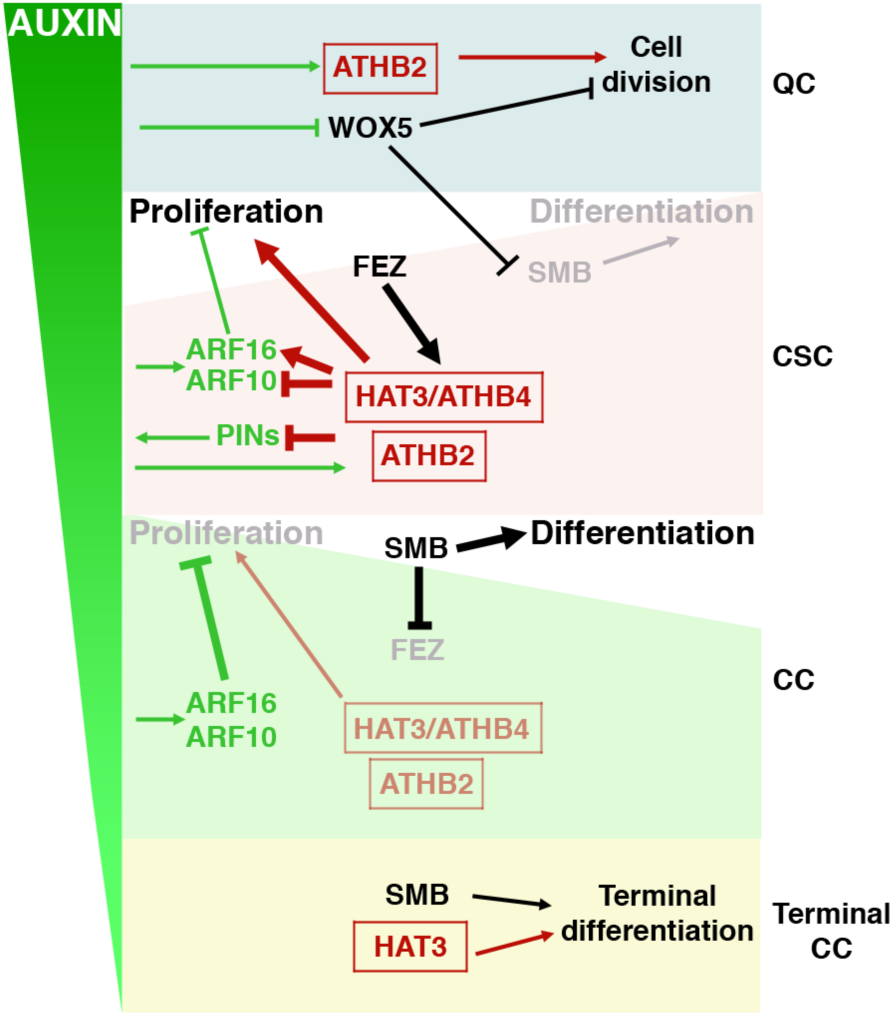
A model for HD-Zip II function in CSCs fate determination. In CSCs, HD-Zip II promote proliferation downstream of FEZ, in part by interfering with auxin-mediated differentiation. On one hand, HD-Zip II affect the expression of PINs to modulate auxin gradients; on the other hand, HD-Zip II control ARF10/ARF16 expression, counteracting their negative role on CSCs divisions. ATHB2 is induced by auxin in a FEZ-dependent manner, ensuring enhanced proliferation to increasing hormone levels. In the QC, increased ATHB2 expression may promote division to replenish differentiating CSCs, independently of WOX5. In CCs, SMB represses FEZ, causing the reduction of HD-Zip II levels. In absence of HD-Zip II, the negative action of auxin on proliferation becomes dominant, leading to CCs differentiation concertedly with SMB. HAT3, which is also expressed in the lower CCs, may contribute to terminal differentiation as inferred from XVE>HAT3 lines.

It was previously shown that HAT3, ATHB4, and ATHB2 redundantly regulate SAM activity and maintenance (Turchi et al., 2013), which, together with the findings reported here, may lead to the idea that HD-Zip II TFs function is to promote proliferation. However, in leaf primordia different members of the HD-Zip II family inhibit the proliferation of mesophyll cells (Carabelli et al., 2018; Challa et al., 2019; Ciarbelli et al., 2008). In particular, upon prolonged exposure to shade, ATHB2 and ATHB4 were shown to slow down cell divisions, leading to an exit from the cell cycle, and eventually resulting in mesophyll cell differentiation (Carabelli et al., 2018). Thus, HD-Zip II TFs’ role in cell fate determination seems to be context dependent: even in the proximal RAM, the induction of XVE>HAT3 or XVE>ATHB4 also promotes the differentiation of proximal epidermal cells into terminal CC, as indicated by the accumulation of starch granules (Figure S8). A number of hypotheses may help explain context-dependent functions of HD-Zip II in cell fate determination. For instance, differences in specific HD-Zip II family members expression, due to environmental and/or hormonal cues (Ciarbelli et al., 2008; Köllmer et al., 2011; Tan et al., 2018; Tan et al., 2021; Zhang et al., 2014), may shift their relative abundance at the tissue level thus altering their ability to form homo-or hetero-dimers with different DNA-binding specificity. Alternatively, tissue-specific differences in chromatin accessibility and/or in the availability of interacting partners may shift HD-Zip II activity towards their non-DNA-binding function as transcriptional co-factors (Gallemí et al., 2017; Zheng et al., 2019). All these hypotheses must be taken into account in future research aiming to understand the role of HD-Zip II in the regulation of cell fate determination.

## Materials and Methods

### Plant lines and growth conditions

Wild type used was Arabidopsis thaliana (L.) Heynh var. Columbia (Col-0). Insertional *HD-Zip* lines used were: *athb2-2* (Salk_006502), *hat3-3* (Salk_014055), *athb2-3 (Turchi et al., 2013)*, *athb4-1* (Salk_104843; Sorin et al., 2009). Multiple *HD-Zip II* mutants used were: *hat3-3 athb4-1*, *hat3-3 athb4-3*, *hat3-2 athb4-1*, *hat3-3 athb2-3*, *athb4-1 athb2-3*, *hat3-3 athb4-1 athb2-1*, *hat3-3 athb4-1 athb2-3*, *hat3-3 athb4-1 athb2-2*, *hat3-3 athb4-1 ATHB2::ATHB2:GUS* (Turchi et al., 2013). Other lines utilized were: *pin3-4* (NASC N9363; Friml et al., 2002a), *pin4-3* (NASC N9368; Friml et al., 2002b), *pin7-2* (NASC N9366; Friml et al., 2003), *wox5-1* (Sarkar et al., 2007), *fez-2* (NASC N678269; Willemsen et al., 2008), *smb-3* (NASC N657070; Willemsen et al., 2008), *arf10-2 arf16-2* (Wang et al., 2005). Marker lines used were: *DR5rev::GFP* (Friml et al., 2003), J2341 (Sabatini et al., 2003), *TAA1::GFP:TAA1* (Stepanova et al., 2008), *HAT3::HAT3:GFP* (Turchi et al., 2013), WOX5::GFP (Sarkar et al., 2007), *FEZ::FEZ:GFP* (Willemsen et al., 2008), pARF10::n3GFP, pARF16::n3GFP (Rademacher et al., 2011), *PIN3::PIN3:GFP*, *PIN4::PIN4:GFP*, *PIN7::PIN7:GFP* (Vieten et al., 2005), DR5rev::GFP (Friml et al., 2003). Plants were vertically grown for 5 days in white light as previously described (Steindler et al., 1999). To analyze the effect of the chemicals, seedlings were grown for 4 days in control medium and then transferred on agar plates supplemented with the appropriate drug for 1 day. Concentrations were as follows: DEX, 10μM; NAA, 5 μM (1 μM for ATHB2:GFP experiments). For NAA, β-estradiol, and NAA + β-estradiol experiments the treatments were as follows: DMSO 2 d; 5 μM NAA 1 d; 10 μM β-estradiol 2 d; β-estradiol + NAA, 10 μM β-estradiol 1d and 10 μM β-estradiol + 5 μM NAA 1d. For auxinole experiments, seedlings were germinated in control medium or in media supplemented with 5 μM auxinole. Equal amounts of DMSO (EtOH, for DEX experiments) were used as control.

### Genetic analysis

The following multiple mutants were generated by crossing: *hat3-3 athb4-1 athb2-3 pin3-4*, *hat3-3 athb4-1 athb2-3 pin4-3*, *hat3-3 athb4-1 athb2-3 pin7-2*, *hat3-3 athb4-1 athb2-3 wox5-1*, *hat3-3 athb4-1 athb2-3 fez-2*, *hat3-3 athb4-1 athb2-3 smb-3*, *hat3-3 athb4-1 athb2-3 arf10 arf16.* All the multiple mutants were selected in F2 by phenotyping and/or PCR genotyping and re-analyzed in F3 (see Table S6 for primer details). Markers J2341, *DR5rev::GFP*, *PIN3::PIN3:GFP*, *PIN4::PIN4:GFP*, *PIN7::PIN7:GFP*, *TAA1::GFP:TAA1*, *WOX5::GFP*, *FEZ::FEZ:GFP* were introduced in *hat3-3 athb4-1 athb2-3* by crossing. *hat3-3 athb4-1 athb2-3 was* selected by phenotyping and genotyping. Homozygosity of all the reporters was determined by unanimous GUS and GFP signal (n≥30).

### Gene constructs and transformation

The following DNA constructs were generated: *HAT3::HAT3:YFP*, *ATHB2::ATHB2:GFP*, *ATHB4::ATHB4:GFP*, *35S::HAT3:GR*, *XVE>ATHB4*, *XVE>HAT3*. Primers used to generate all the constructs are listed in Table S4. Col-0 and *hat3-3 athb4-1* plants were transformed as previously described (Steindler et al., 1999). *HAT3::HAT3:YFP*, *ATHB2::ATHB2:GFP*, were also introduced in different mutant backgrounds by crossing. Mutant backgrounds were selected by genotyping; homozygosity of the DNA constructs was verified by antibiotic resistance of T3 seedlings. *35S::HAT3:GR* was also introduced in *fez-2* mutant.

### Phenotypic analysis and microscopy

Lugol’s staining was performed as previously described (Willemsen et al., 1998). For differential interference contrast (DIC) analysis, roots were cleared in chloral hydrate (Weigel and Glazebrook, 2002), mounted on microscope slides and viewed under an Axioskop 2 plus binocular microscope (Zeiss, Germany). Images were taken with a Axiocam ERc5s Zeiss camera. Means were compared using t-test analysis. Phenotypic distributions were compared using a contingency table followed by Fisher’s exact (test.http://graphpad.com/quickcalcs/contingency1.cfm). For all the experiments, at least 40 samples for each genotype were analyzed.

### F-ara-EdU staining

F-ara-EdU staining was performed as described by (Bennett et al., 2014). 4-day-old seedlings were transferred to plates containing 3 μM F-ara-EdU [(2ʹS)-2ʹ-deoxy-2ʹ-fluoro-5-ethynyluridine] for 1 day and were then fixed in 4% paraformaldehyde. Detection of F-ara-EdU was performed using a Click-iT EdU Alexa Fluor 555 Imaging Kit (Life Technologies) according to the manufacturer’s instructions. Seedlings were counterstained with 0.1 μg/ml DAPI (4,6-diamidino-2-phenylindole) and F-ara-EdU incorporation was assessed by confocal microscopy.

### Confocal microscopy

Confocal microscopy analyses on roots expressing reporter genes were performed on an Inverted Z.1 microscope (Zeiss, Germany) equipped with a Zeiss LSM 700 spectral confocal laser scanning unit (Zeiss, Germany). Samples were excited with a 488 nm, 10mW solid laser with emission at 492-539 nm for Green Fluorescent Protein (GFP) detection, and with emission at 492-549 nm for Yellow Fluorescent Protein (YFP) detection. The seedlings were grown for 5 days and analysed. Counterstaining of cell walls was achieved by mounting seedling roots in 10 μM propidium iodide (PI). For F-ara-EdU imaging, excitation was performed using 405 nm and 555 nm lasers; Alexa Fluor 555 fluorescence was detected above 555 nm and DAPI fluorescence below 500 nm.

## Acknowledgments

With this work, we would like to remember our friend, colleague, and woman in science Ida Ruberti who sadly passed away during the middle stage of this project, for her instrumental role in the conceptualization of the project and the many great discussions of the results. We thank Sabrina Sabatini for providing the J234 marker line, Ben Scheres for the FEZ::FEZ:GFP line, Xiao-Ya Chen for the *arf10-2 arf16-2* line, and Laila Moubayidin for discussions and helpful comments on the manuscript.

## Author contribution

Conceptualization: G.M., I.R.; Methodology: M.P., G.S., A.A., L.T., V.R. M.S., G.M., I.R.; Formal analysis: M.P., G.S., A.A., M.S., G.M., I.R.; Investigation: M.P., G.S., A.A., L.T., V.R. M.S.; Data curation: M.P., G.S., M.S., G.M., I.R.; Writing - original draft: M.S., G.M., I.R.; Writing - review & editing: M.P., G.S., M.S., G.M.; Supervision: G.S., M.S., G.M., I.R.; Funding Acquisition: G.S., G.M., I.R.

## Data availability

All relevant data can be found within the article and its supplementary information.

